# The multidimensional structure of wellbeing: genetic evidence from a multivariate twin study including the Mental Health Continuum

**DOI:** 10.64898/2026.03.27.714768

**Authors:** Natalia Azcona-Granada, Anne Geijsen, Lianne de Vries, Dirk Pelt, Meike Bartels

## Abstract

Wellbeing is commonly defined as the combination of feeling good and functioning well and typically conceptualized as two related but distinct components. Hedonic wellbeing emphasizes pleasure, happiness, and life satisfaction, while eudaimonic wellbeing focuses on meaning, personal growth, flourishing, and the realization of one’s potential. The Mental Health Continuum-Short Form was developed as a comprehensive measure of wellbeing and includes three subscales assessing emotional, social, and psychological wellbeing. Although the Mental Health Continuum total score is often interpreted as an indicator of overall wellbeing, the underlying genetic structure of its three subscales and its genetic overlap with other commonly used wellbeing measures remains unclear.

Using data from 5,212 individuals from the Netherlands Twin Register (72% female, mean age 36.4), we fitted multivariate twin models to examine the genetic architecture of the Mental Health Continuum and its associations with other wellbeing measures (quality of life, life satisfaction, subjective happiness, and flourishing). Results indicate that, at the genetic level, the Mental Health Continuum is best explained by its three distinct subscales rather than by a latent factor. When considering the Mental Health Continuum together with the other wellbeing measures, we found moderate to high genetic correlations (*r* = 0.52 – 0.83), indicating substantial overlap in the genetics underlying the wellbeing constructs. However, we did not find evidence for a single common genetic factor underlying all constructs. These findings highlight the multidimensional structure of wellbeing, but the moderate to high genetic correlations across measures suggest that it is important to align the level of measurement (phenotypic vs genetic) with the research question.

## Introduction

Wellbeing can be defined as the combination of feeling good and functioning well (OECD, 2013; Organization, 2022). Wellbeing is widely understood as a broad, multidimensional construct encompassing how people feel, how they function personally and socially, and how they evaluate their lives as a whole (Jarden et al., 2023). It is measured through a variety of domain□specific and composite indicators such as life satisfaction, positive affect, meaning in life, social relationships, and health or capability-based indices (OECD, 2024).

Theoretically, wellbeing is often conceptualized in terms of two related but distinct components: hedonic wellbeing, which emphasizes pleasure, happiness, and life satisfaction; and eudaimonic wellbeing, which focuses on meaning, personal growth, flourishing, and fulfilling one’s potential (Ryan & Deci, 2001). Over the past decade, wellbeing research has expanded considerably, producing a wide array of constructs such as life satisfaction, happiness, flourishing, and overall quality of life. Despite the conceptual distinctions made between these constructs, recent findings argue that progressively complex models of wellbeing may promise specificity but often show overlapping structures of wellbeing instead (Helliwell et al., 2026; Sametoğlu et al., 2023). In other words, adding new measures of wellbeing may provide only the illusion of measuring distinct constructs.

Most behavior genetic research has been done on hedonic measures, including quality of life (QoL; (Cantril, 1965)), satisfaction with life (SWLS; (Diener et al., 1985)), and happiness (SHS; (Lyubomirsky & Lepper, 1999)). Often, such measures are interchangeably used to provide information on wellbeing even though they capture partly distinct aspects of wellbeing as indicated by phenotypic correlations ranging from 0.57 to 0.69 in the most recent meta-analysis (Lomas et al., 2026).

However, these aforementioned measures do not include eudaimonic wellbeing. One widely used measure of eudaimonic wellbeing is the Flourishing Scale (Diener et al., 2010), an 8-item instrument assessing purpose, self-esteem, engagement, optimism, and social relationships, providing a broad and unidimensional evaluation of eudaimonic wellbeing and psychological prosperity (Diener et al., 2010).

In contrast to the aforementioned wellbeing measures, focusing on either hedonic or eudaimonic wellbeing, the Mental Health Continuum-Short Form (MHC) arose as a comprehensive measure of both hedonic and eudaimonic wellbeing, assuming to capture “overall” wellbeing (Lamers et al., 2011). The MHC consists of three subscales; emotional, social and psychological wellbeing. Earlier studies have reported phenotypic correlations of the Mental Health Continuum with life satisfaction (*r* = 0.35 – 0.40) (Keyes et al., 2008), and flourishing (*r* = 0.57 – 0.73) (Ploke et al., 2024). Phenotypic correlations with the hedonic measures (life satisfaction) are relatively moderate while the phenotypic correlation with flourishing (eudaimonic measure) is relatively high, suggesting that although the Mental Health Continuum captures wellbeing broadly, it somewhat more reflects eudaimonic wellbeing than hedonic wellbeing (Iasiello et al., 2022; Lamers et al., 2011).

Although phenotypic associations indicate overlap among wellbeing measures, it remains unclear to what extend this overlap reflects shared genetic influences. The etiology of wellbeing arises from the complex interplay of genetic and environmental influences that shape individual differences (Bartels et al., 2022; Bartels & Boomsma, 2009). Twin studies have demonstrated that wellbeing is moderately heritable, with genetic factors explaining approximately 40% of its variance, depending on how it is measured and modeled (Bartels, 2015). Evidence from behavioral genetic studies suggests that commonalities among different wellbeing measures may outweigh their differences (Bartels & Boomsma, 2009; Baselmans & Bartels, 2018; Bjørndal et al., 2023; Pelt et al., 2024; Sametoğlu et al., 2025). Specifically, genetic correlations among wellbeing indicators are consistently found to be high, often higher than their phenotypic correlations, implying that the same underlying genetic factors may influence multiple dimensions of wellbeing (Pelt et al., 2024). Meanwhile, nonshared environmental influences, such as life experiences, social relationships, and stress exposure, influence individual differences in wellbeing throughout life (Kendler et al., 2011; van de Weijer et al., 2022).

This study examines whether the Mental Health Continuum captures unique genetic variance beyond that explained by other wellbeing measures, like happiness, satisfaction with life, and flourishing. We also test whether the Mental Health Continuum is best represented as a single overarching wellbeing factor or by its three correlated subscales: emotional, social, and psychological wellbeing. Using multivariate twin models, we assess the shared and unique genetic contributions to the Mental Health Continuum (total and subscale scores) and its covariance with quality of life, life satisfaction, subjective happiness, and flourishing. Finally, we estimate genetic and environmental correlations among these measures to determine whether the Mental Health Continuum reflects distinct aspects of wellbeing or overlaps with the other constructs. Our research questions are:

- How should wellbeing as assessed by the Mental Health Continuum–Short Form be conceptualized genetically: as a single overarching construct (either as a latent factor or the total score) or as three related but distinct domains of emotional, psychological, and social wellbeing?
- To what extent does wellbeing captured by the Mental Health Continuum–Short Form overlap with other commonly used wellbeing measures, including quality of life, life satisfaction, subjective happiness, and flourishing, at the level of underlying genetic and environmental influences?

## Materials and methods

### Participants

Participants included in this study are part of the Netherlands Twin Register (NTR), a national register of twins and their families with longitudinal data collection on mental health, personality, lifestyle, demographics (Ligthart, van Beijsterveldt, et al., 2019). The inclusion criteria for being part of the NTR are to be Dutch-literate and a twin or a relative of a twin. This study included data from the survey completed in 2019 – 2022. Being part of a national prospective cohort study, Netherlands Twin Register data cannot be made publicly available for privacy reasons, but they are available for researchers via the data access procedure (https://ntrdata.nl/).

All procedures performed in studies involving human participants were in accordance with the ethical standards of the institutional and/or national research committee and with the 1964 Helsinki declaration. Data collection was approved by the Central Ethics Committee on Research Involving Human Subjects of the University Medical Centers Amsterdam (Ethical approval number: 2018-389 (25-07-2018)). Informed consent was obtained from all individual participants included in the study.

Data were available for 5,212 individuals, including 1,024 complete twin pairs, from 4,188 families. Twins were either identical twins (monozygotic twins, MZ, who share nearly all the same DNA, N=2810) or fraternal twins (dizygotic twins, DZ, who share on average half of their DNA, N=2402). The mean age for the twins was 36.36 [standard deviation (*SD*) 15.08, range 16-92 years].

Zygosity was determined either by genotyping (55.5%) or from self- and parental report answers to survey questions on physical resemblance. DNA and survey zygosity agreement reached more than 96% (Ligthart, Van Beijsterveldt, et al., 2019).

### Measures

Wellbeing was assessed using the following measures:

Quality of Life (QoL) was assessed with the Cantril ladder, which requires participants to indicate the step of the ladder at which they place their lives in general on a 10-point scale (10 being the best possible life, and 1 the worst possible life) (Cantril, 1965).

Life satisfaction (LS) was assessed with the satisfaction with life scale (SWLS) (Diener et al., 1985). The SWLS consists of five items answered on a seven-point scale ranging from 1 = “strongly disagree” to 7 = “strongly agree.” An example item is: “I am satisfied with my life”. A total score was calculated by summing up all the items, and Cronbach’s alpha was 0.87.

Subjective Happiness (SH) was assessed with the Subjective Happiness Scale (SHS) (Lyubomirsky & Lepper, 1999). It consists of four items which are rated on a seven-point scale ranging from 1 = “strongly disagree” to 7 = “strongly agree.” An example item is “On the whole, I am a happy person”. A total score was calculated by summing up all the items, and Cronbach’s alpha was 0.89.

Flourishing (FL) was assessed using the Short Flourishing Scale (Diener et al., 2010). It contains 8 items that are rated from 1 to 7 using a Likert scale. 1 resembles “strongly disagree” and 7 resembles “strongly agree”. An example of an item is: “I am competent and capable in the activities that are important to me”. A total score was calculated by summing up all the items, and Cronbach’s alpha was 0.90.

The Mental Health Continuum-Short Form is a self-report questionnaire that assessed positive mental health. The questionnaire consists of 14 items measuring positive mental health (Keyes, 2002). These items measure emotional (3 items), social (5 items) and psychological wellbeing (6 items) and are scored on a 6-point scale from never (0) to every day (5). Some example questions are: During the past month, how often did you feel “… happy?”; “… that you belonged to a community?”; “… that people are basically good?”. The three subscales were used individually by taking the sum of the items for each scale. In addition, the total score was calculated by averaging all the items following the guidelines (Keyes, 2002). We refer to this measure as MHC and to the subscales as emotional MHC (EMHC), social MHC (SMHC), and psychological MHC (PMHC), respectively. Cronbach’s alpha was 0.92 for MHC, 0.87 for EMHC, 0.8 for SMHC, and 0.87 for PMHC.

### Statistical Analyses

Descriptive analyses were done in R v.4.3.2. (*R: The R Project for Statistical Computing*, 2021). We performed all modeling and testing in RStudio, notably with the packages “OpenMx” v2.19.8 (S. M. Boker et al., 2025), "psych" v2.1.6 (Revelle, 2025), "ggplot2" v3.3.5 (Wickham et al., 2025), "foreign" 0.8-81 (R Core Team, 2023).

#### Descriptives

First, we estimated phenotypic correlations between the Mental Health Continuum and the other wellbeing measures in a saturated model. Using the saturated model, i.e., a model in which all means, variances, and covariances are estimated freely, we tested the assumptions of the classical twin model (equal means and variances across twin order and zygosity). We estimated the means, variances, covariances, and twin correlations (MZ and DZ) of the individual wellbeing measures, i.e., quality of life, life satisfaction, subjective happiness, flourishing, Mental Health Continuum subscales. Age and sex were included as covariates in the saturated model and we tested the effect of age and sex on the means of the wellbeing measures.

#### Multivariate twin modelling

Second, we ran different multivariate twin models on the wellbeing measures, including quality of life, life satisfaction, subjective happiness, flourishing, and Mental Health Continuum to estimate to what extent wellbeing captured by the Mental Health Continuum-Short Form overlaps with the other measures estimating the genetic and environmental variances and covariances, and calculating genetic and environmental correlations.

The contribution of genetic and non-genetic influences in the classical twin models is based on the different degrees of genetic relatedness between monozygotic (MZ) and dizygotic (DZ) twins. MZ twins are (nearly) 100% genetically similar, while DZ twins share ∼50% of their segregating genes (Beck et al., 2021). Because both types of twins are born at the same time and grow up in the same household, MZ and DZ twins share features of their environment and experiences – labelled the shared environment (C), which might affect their trait resemblance beyond genetic similarity. Unique environmental factors and experiences (E) cause differences within MZ and DZ pairs and include all influences associated with “unique” environments (aspects of the environment that differ among siblings), and all forms of error and random noise. E factors are correlated zero by definition in MZ and DZ pairs. When there is a higher resemblance between MZ twins than between DZ twins for a trait, this indicates an influence of genetic factors. Twin resemblance can be a function of additive genetic influences (A), or an influence from interactions between alleles at the same locus (dominance (D)) or between alleles at different loci (epistasis) (Neale & Cardon, 2013).

Genetic structural equation modelling (Bruins et al., 2023; Neale & Cardon, 2013) to estimate the contribution of genetic and environmental influences was performed in OpenMx (S. Boker et al., 2011). Decomposition of the *variance* indicates how much of the variation within the traits is accounted for by genetic and environmental factors. The *covariance* decomposition answers the question of how much of the phenotypic correlation between the traits is accounted for by genetic and environmental factors. The proportion of covariance explained by genetic factors is called the bivariate heritability (Eaves & Gale, 1974; Martin & Eaves, 1977).

Additionally, in a multivariate model, the *genetic and environmental correlations* between the traits can be calculated. The genetic correlation reflects the extent to which the genetic factors underlying each trait overlap. Similarly, the environmental correlation reflects the overlap of the environmental factors underlying the traits. A genetic (or environmental) correlation of 1 indicates a perfect overlap of the genetic (or environmental) factors, indicating that the influences on both traits are identical. In contrast, a correlation of 0 indicates no overlap and thus independent genetic (or environmental) factors.

In classical twin designs, an ACE model is fitted when the dizygotic twin correlation is at least half the monozygotic correlation (suggesting shared environmental effects), whereas an ADE model is preferred when the dizygotic correlation is less than half the monozygotic correlation (suggesting non-additive genetic effects), after which nested submodels are compared using likelihood-ratio tests and information criteria. Based on the pattern of twin correlations obtained from the saturated model 1) We test a ‘no shared environment’ AE model by constraining the common environmental component (C) to zero in an ACE model. If the starting model is ADE, we evaluate a ‘no dominance’ AE model by fixing the dominance genetic component (D) to zero, leaving only additive genetic and unique environmental effects. 2) If the starting model is ACE, we test a ‘no additive genetics’ CE model by dropping the A component, so that variance is explained only by shared and unique environmental influences. 3) We consider an ‘E-only’ model in which both the genetic component (A) and the shared or dominance component (C or D, depending on the starting model) are constrained to zero, such that all variance is attributed to unique environmental effects and measurement error.

The fit of the different models (both for the saturated as well as the genetic models) was compared by means of the log-likelihood ratio test (LRT). The difference in minus two times the log-likelihood (−2LL) between two nested models has a χ 2 distribution with the degrees of freedom (df) equaling the difference in df between the two models. If a p value higher than 0.05 was obtained from the χ 2 test, the fit of the constrained model was considered not significantly worse than the fit of the more complex model. In this case, the constrained model was kept as the most parsimonious and best-fitting model.

#### Comparing different approaches to modelling MHC

To analyze the optimal structure of the Mental Health Continuum questionnaire, three multivariate twin models (and submodels) were applied, including quality of life, life satisfaction, subjective happiness, flourishing, and Mental Health Continuum in all models.

1. In the 7-variable ACE model, quality of life, life satisfaction, subjective happiness, flourishing, and the three subscales of the Mental Health Continuum, i.e., emotional, social and psychological, were included. This model implies that genetically, the Mental Health Continuum should be treated as three different but correlated subscales.
2. In the 5-variable ACE model, quality of life, life satisfaction, subjective happiness, flourishing, and the Mental Health Continuum total score, calculated following the guidelines of the Mental Health Continuum (Keyes, 2002), were included. This model implies that, genetically, the Mental Health Continuum should be treated as one wellbeing score, as implied by the questionnaire manual.
3. In the common pathway ACE model, the Mental Health Continuum was included as a latent factor calculated from the three aforementioned subscales. Besides the Mental Health Continuum latent factor, quality of life, life satisfaction, subjective happiness, and flourishing, were included. In common pathway models, the genetic and environmental effects are mediated by one or more latent phenotypic factors, which capture the phenotypic overlap (i.e., the shared variance) among its indicators. In addition to variance components that capture the variance common to multiple traits, specific variance components for each trait can be modeled. Thus, genetic and environmental influences cause individual differences in the latent phenotype, in turn influencing the observed indicators. This model implies that, genetically, the Mental Health Continuum subscales are influenced by a common latent factor, albeit to different degrees.

### Best-fitting models/model comparison

To obtain the best fitting model for the Mental Health Continuum, likelihood tests were performed between the nested models, i.e., the model considering the 3 subscales separately (7-variable ACE model) and the model considering a latent factor (common pathway ACE model). However, the model considering the Mental Health Continuum as an average score (5-variable ACE model) cannot directly be compared with the other models as it is not nested under these models. The 5-variable ACE model is based on the assumption that there is an underlying factor influencing the three Mental Health Continuum subscales (emotional, social and psychological) and therefore any conclusion regarding the common pathway model was extended to the model considering the average score (5-variable ACE model).

### Genetic and environmental factors of wellbeing measures

Once established which model fits the Mental Health Continuum data best, we aimed to test whether all the wellbeing measures are all influenced by a single common genetic factor, as previously found for quality of life, life satisfaction, and subjective happiness (Bartels & Boomsma, 2009). For this aim, an independent pathway model was estimated including quality of life, life satisfaction, subjective happiness, flourishing and the best-fitting structure for the Mental Health Continuum (one overall scale or three subscales). In independent pathway models, A, C/D, and E are conceptualized as one or more (orthogonal or correlated) latent global factor(s) directly influencing the phenotypes. In addition to variance components that capture the variance common to multiple traits, specific variance components for each trait can be modeled. These residual factors capture the genetic and environmental variance in each phenotype that remains after common genetic and environmental effects are accounted for. For an overview of the models, see Figure 1.

**Fig. 1.**
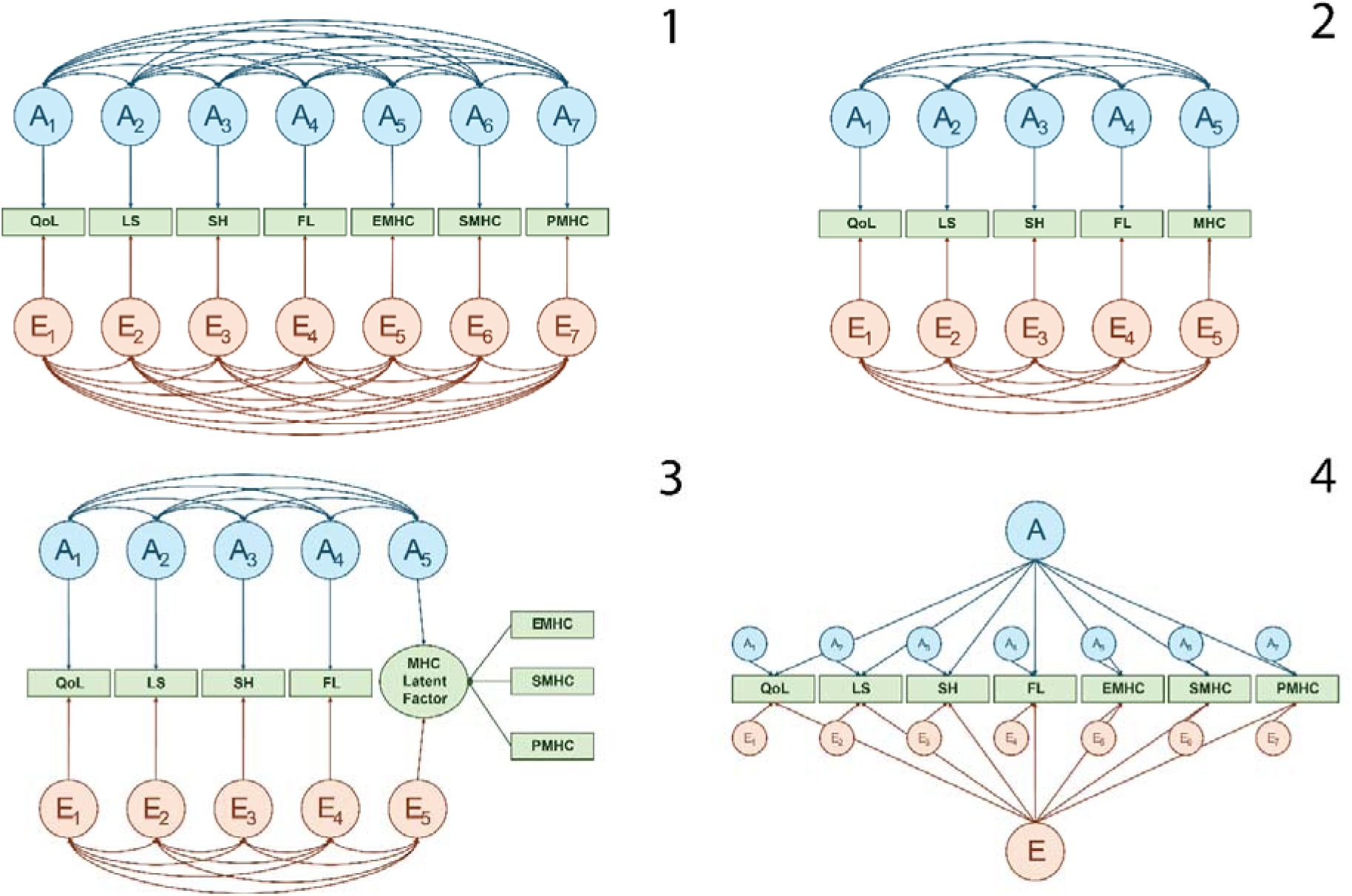
Overview of the AE models fitted. Overview of the AE models fitted. For simplification, C/D component was not represented. Models 1, 2 and 3 were fitted to understand the genetic and environmental structure of the Mental Health Continuum data. Model 1 is the 7-variable AE model. Model 2 is the 5-variable AE model. Model 3 is the common pathway AE model. Model 4 is an independent pathway model, in which we added the three subscales for the MHC, but it could also be the single overall score. In green, the wellbeing phenotypes. In blue, the genetic components, where a straight arrow indicates the unique genetic contribution to a phenotype and the curved arrows indicate the genetic correlation among the wellbeing phenotypes. In orange, the non-shared environmental components, where a straight arrow indicates the unique non-shared contribution to a phenotype, and the curved arrows indicate the environmental correlation among the wellbeing phenotypes. QoL: Quality of Life; LS: Life Satisfaction; SH: Subjective Happiness; FL: Flourishing; EMHC: Emotional Mental Health Continuum, SMHC: Social Mental Health Continuum, PMHC: Psychological Mental Health Continuum.

**Fig. 2.**
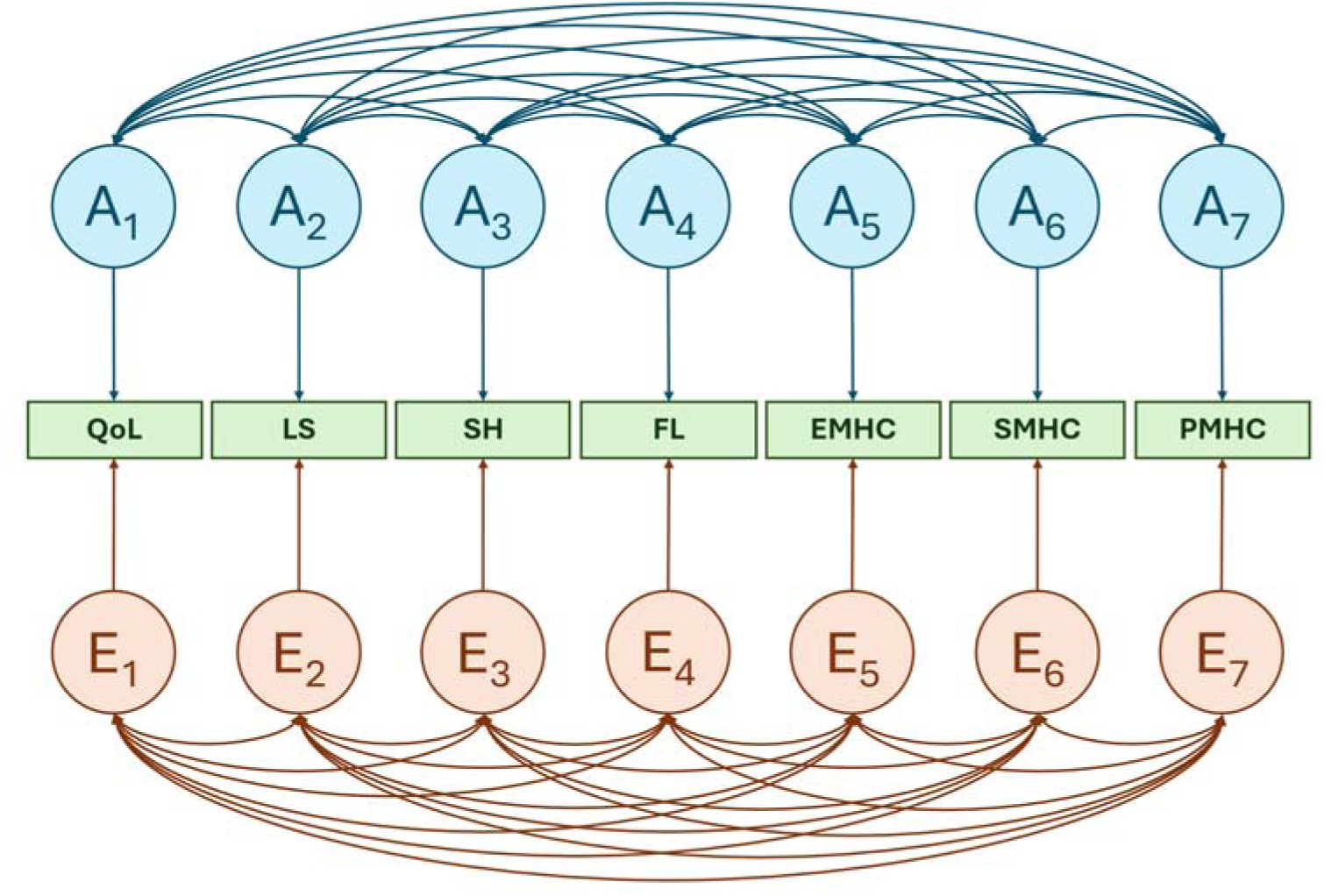
Best-fitting multivariate twin model. Best-fitting multivariate twin model. In green, the wellbeing phenotypes. In blue, the genetic components, where a straight arrow indicates the unique genetic contribution to a phenotype and the curved arrows indicate the genetic correlation among the wellbeing phenotypes. In orange, the non-shared environmental components, where a straight arrow indicates the unique non-shared contribution to a phenotype, and the curved arrows indicate the environmental correlation among the wellbeing phenotypes. QoL: Quality of Life; LS: Life Satisfaction; SH: Subjective Happiness; FL: Flourishing; EMHC: Emotional Mental Health Continuum, SMHC: Social Mental Health Continuum, PMHC: Psychological Mental Health Continuum.

## Results

### Descriptive statistics and correlations

Phenotypic correlations from the saturated model are presented in Table 1. These correlations ranged between 0.50 – 0.76, indicating a moderate to large overlap between the seven measures of wellbeing.

**Table 1.** Phenotypic correlations constrained across MZ and DZ.

|  | QoL | LS | SH | FL | EMHC | SMHC | PMHC | MHC |
| --- | --- | --- | --- | --- | --- | --- | --- | --- |
| QoL | 1 | 0.76 | 0.74 | 0.69 | 0.59 | 0.37 | 0.49 | 0.52 |
| LS |  | 1 | 0.75 | 0.71 | 0.56 | 0.36 | 0.48 | 0.50 |
| SH |  |  | 1 | 0.72 | 0.63 | 0.39 | 0.52 | 0.56 |
| FL |  |  |  | 1 | 0.59 | 0.50 | 0.62 | 0.64 |
| EMHC |  |  |  |  | 1 | 0.57 | 0.68 | 0.79 |
| SMHC |  |  |  |  |  | 1 | 0.72 | 0.90 |
| PMHC |  |  |  |  |  |  | 1 | 0.93 |
| MHC |  |  |  |  |  |  |  | 1 |

Note. QoL: Quality of Life; LS: Life Satisfaction; SH: Subjective Happiness; FL: Flourishing; EMHC: Emotional Mental Health Continuum, SMHC: Social Mental Health Continuum, PMHC: Psychological Mental Health Continuum.

The joint effects of age and sex on the means of the wellbeing measures were significant (*P_FDR_* = 0.00 for both tests), see Supplementary Table 1. When the effect of age or sex was constrained to 0 for the different wellbeing measures independently, only the effect of sex on subjective happiness, flourishing and emotional Mental Health Continuum (*P_FDR_* = 0.29, 0.44, 0.92, respectively) could be constrained. Men report on average higher quality of life (β = −0.11, SE = 0.03), and life satisfaction (β = −0.06, SE = 0.03), while women report on average higher social wellbeing (β = 0.09, SE = 0.03), and psychological wellbeing (β = 0.09, SE = 0.03). Quality of life (β = 0.61, SE = 0.10), life satisfaction (β = 0.53, SE = 0.09), subjective happiness (β = 0.55, SE = 0.09), flourishing (β = 0.48, SE = 0.09), emotional wellbeing (β = 0.38, SE = 0.09), social wellbeing (β = 0.28, SE = 0.09), and psychological wellbeing (β = 0.27, SE = 0.10) increased with age. For details regarding the saturated model, see Supplementary Table 2.

### Twin correlations

Cross-twin and cross-twin cross-trait correlations for the wellbeing measures are presented in Table 2. Cross-twin correlations indicate genetic influences on all measures, since the MZ correlations are higher than the DZ correlations. Effects of common environmental factors are to be expected, since the MZ correlations are smaller than 2*DZ correlations for quality of life and life satisfaction. The MZ-DZ differences in the cross-trait correlations, represented in Table 2 by the off-diagonal elements, indicate that the phenotypic correlations between the measures of wellbeing are at least partly accounted for by genetic influences.

**Table 2.**
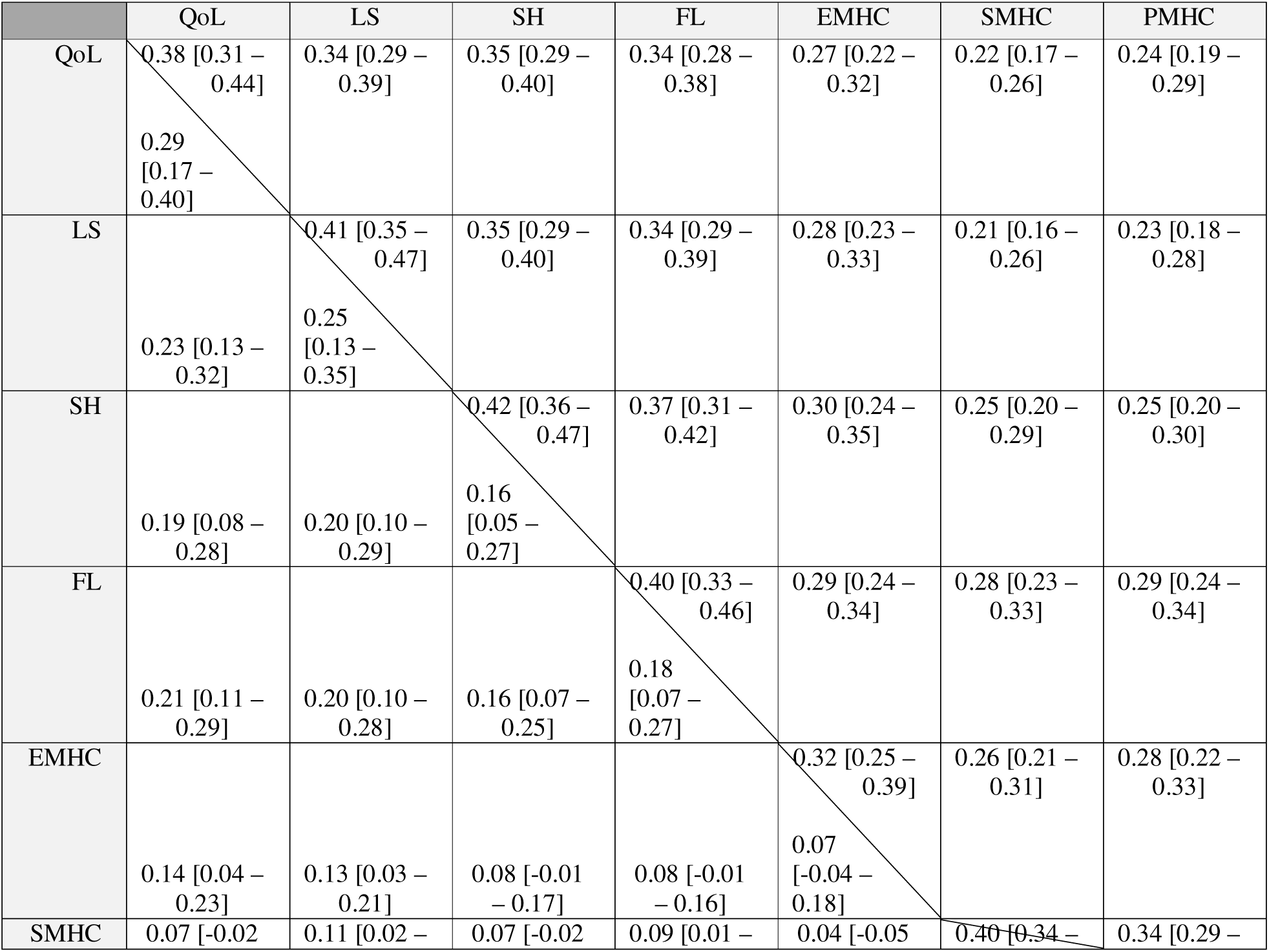
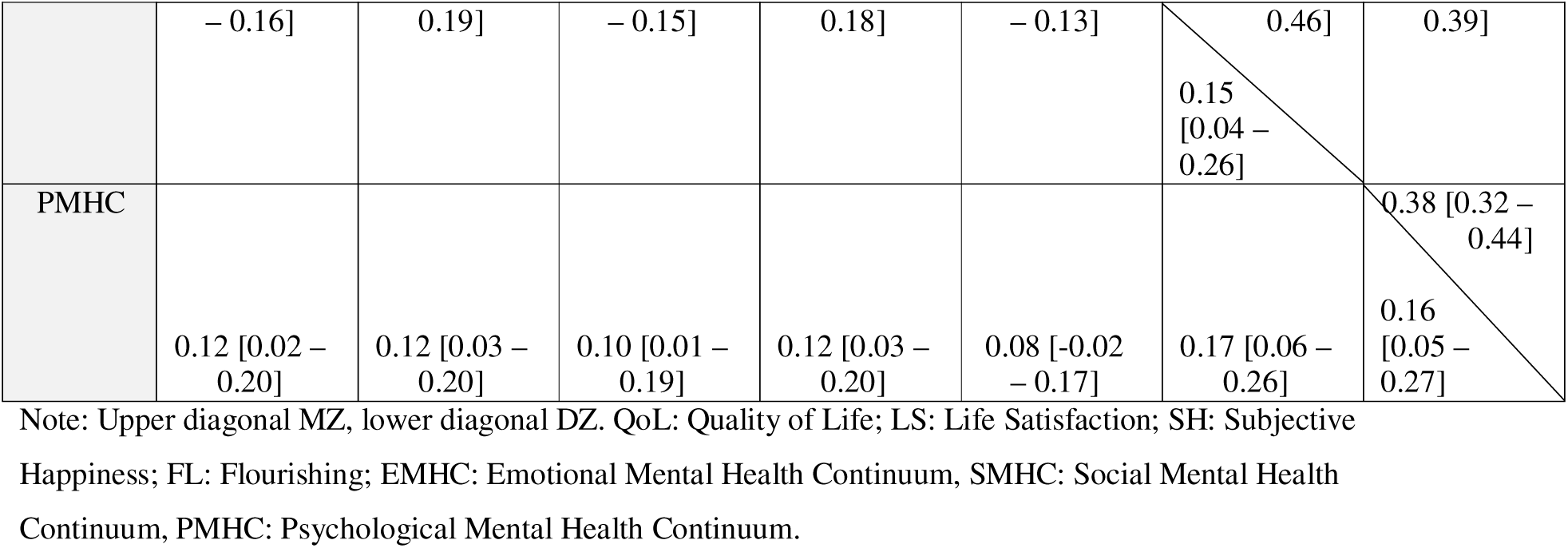
Diagonal cross-twin correlation. Off-diagonal cross-twin cross-trait correlation.

### Comparison of the different multivariate models for MHC

For all three multivariate models, 7-variable ACE model, 5-variable ACE model, and common pathway ACE model, the best fitting model was an AE model (see Supplementary Table 2).

The comparison between-AE models (7-variable AE model vs common pathway AE model) showed that the 7-variable AE model was the best fitting model (See Table 3). Comparisons with the 5-variable AE model was not possible due to the different fit function of the models. However, the 5-variable AE model is based on the assumption that there is a common underlying factor influencing all the three MHC subscales (emotional, social and psychological). Given that the common pathway model did not fit well, creating a sum score of the three subscales (the approach in 5-variable AE model) as a proxy for the general MHC wellbeing factor is not supported by the data. Therefore, the 5-variable AE model was discarded.

**Table 3.**
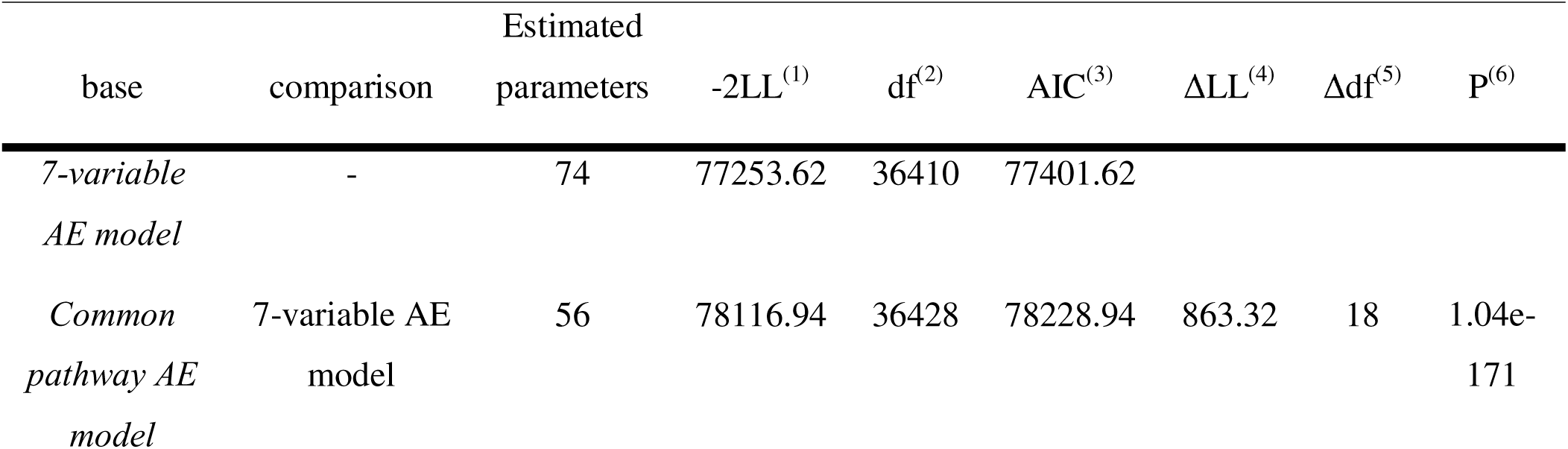
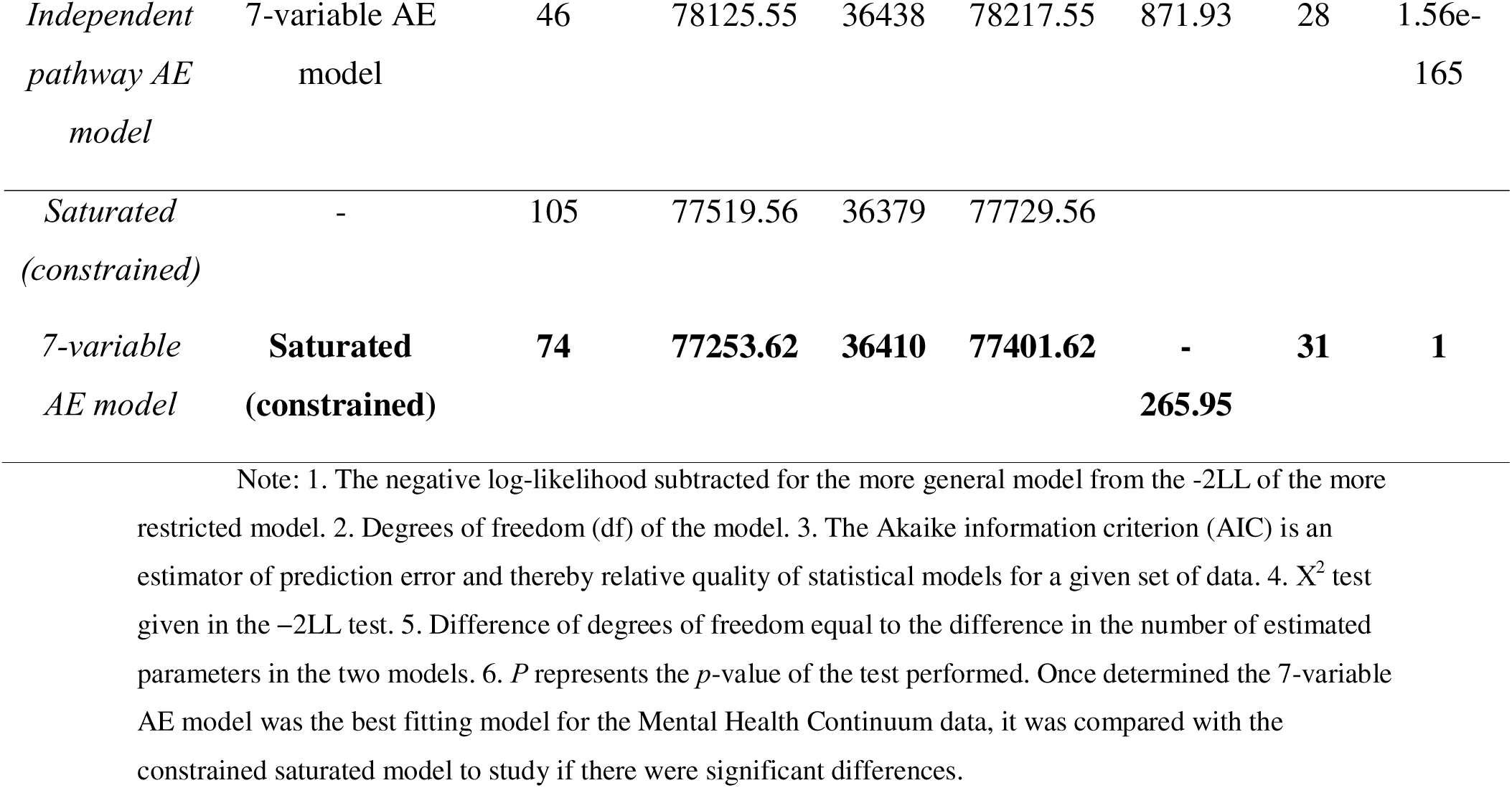
Comparison of models fitted.

#### Understanding the genetic structure of wellbeing

Continuing with the 7-variable AE model, to investigate the common and unique genetic and environmental structure of the wellbeing measures, we fitted an independent pathway model. The comparison between the 7-variable AE model and the independent pathway model showed that there is no evidence for a common genetic and/or environmental factor underlying all the wellbeing measures, as indicated by the worse fit of the independent pathway model (Table 3). For a detailed results of all the comparisons within-model (e.g., ACE vs AE vs E) we performed, see Supplementary Table 3.

#### Standardized variances

Based on the 7-variable AE model, the heritability, i.e., standardized genetic variance, of the Mental Health Continuum is 0.31 [95% CI = 0.24 - 0.37] for emotional wellbeing, 0.40 [95% CI = 0.34 - 0.45] for social wellbeing and 0.37 [95% CI = 0.31 - 0.43] for psychological wellbeing. The bivariate heritability, standardized genetic covariance, of the Mental Health Continuum and the other four wellbeing measures ranges from 0.46 [95% CI = 0.37 - 0.54] for the correlation of the emotional subscale with subjective happiness to 0.63 [95% CI = 0.52 - 0.73] for the correlation of social wellbeing with subjective happiness, indicating that around half of the phenotypic association between the Mental Health Continuum scores and the other measures can be explained by genetic factors (see Table 4). For the unstandardized variances, see Supplementary Table 4.

**Table 4.**
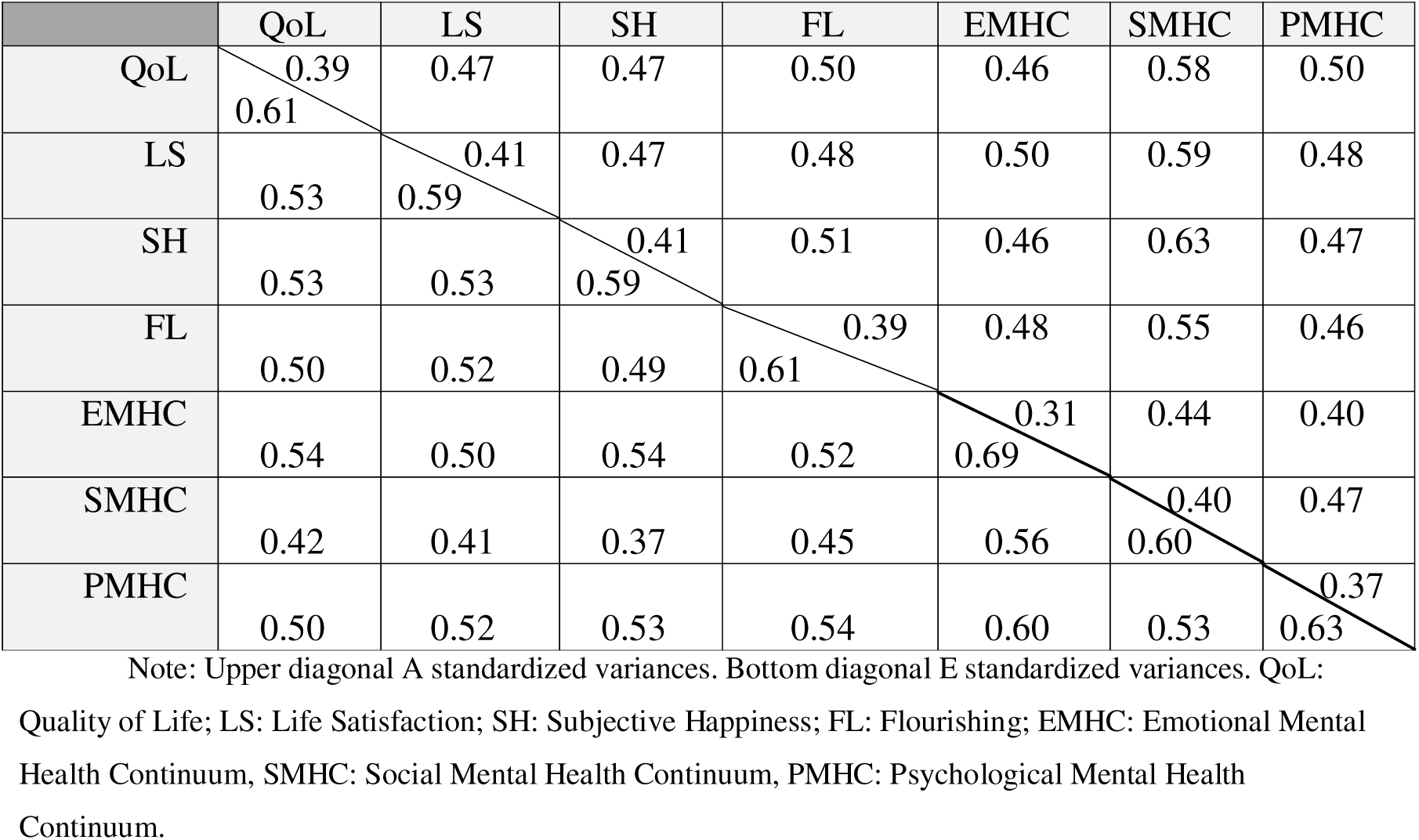
Standardized variances.

The phenotypic correlations between each of the three subscales of Mental Health Continuum and the four other wellbeing measures and the proportions that were accounted for by genetic (i.e., the bivariate heritability), and non-shared environmental influences are presented in Figure 3.

**Fig. 3.**
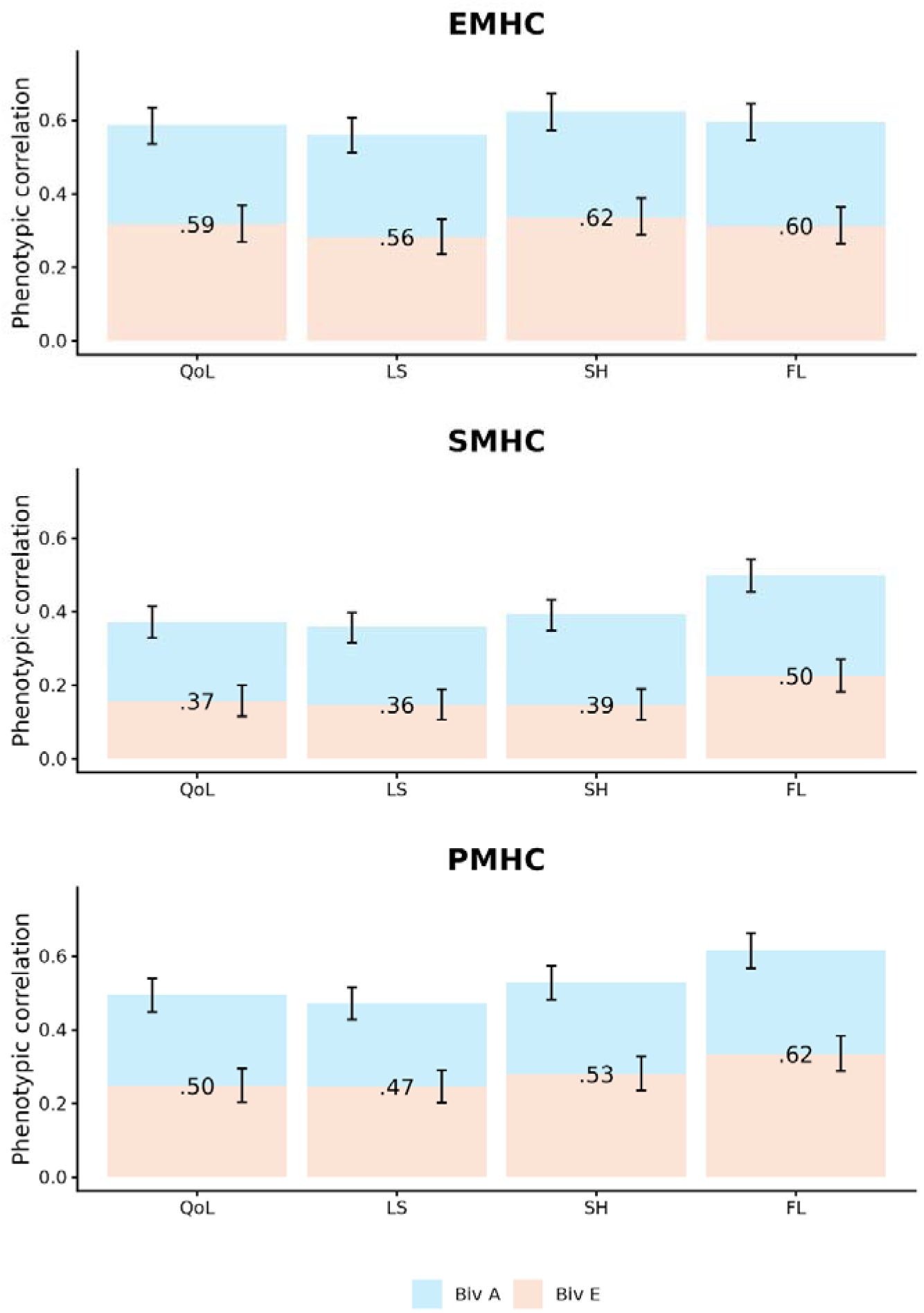
Phenotypic correlations between the three subscales of the Mental Health Continuum and the traits: Phenotypic correlations between the three subscales of the Mental Health Continuum and the traits: quality of life (QoL), Life Satisfaction (LS), Subjective Happiness (SH), and Flourishing (FL), with the proportions that are accounted for by additive genetic (Biv A) and non-shared environmental influences (Biv E). The error bars indicate the 95% confidence intervals. EMHC: Emotional Mental Health Continuum; SMHC: Social Mental Health Continuum: PMHC: Psychological Mental Health Continuum.

#### Genetic and environmental correlations

The genetic correlations between the Mental Health Continuum subscales amongst each other ranged from 0.73 [95% CI = 0.64-0.81] to 0.89 [95% CI = 0.83-0.94]. The genetic correlations between the Mental Health Continuum subscales and quality of life, life satisfaction, subjective happiness and flourishing ranged between 0.52 [95% CI = 0.43-0.62] and 0.83 [95% CI = 0.75-0.91] (Figure 4). At the same time, genetic correlations between quality of life, life satisfaction, subjective happiness and flourishing ranged from 0.85 [95% CI = 0.80-0.90] to 0.92 [95% CI = 0.87-0.97]. This indicates that a substantial part of the genes between these measures overlap.

**Fig. 4.**
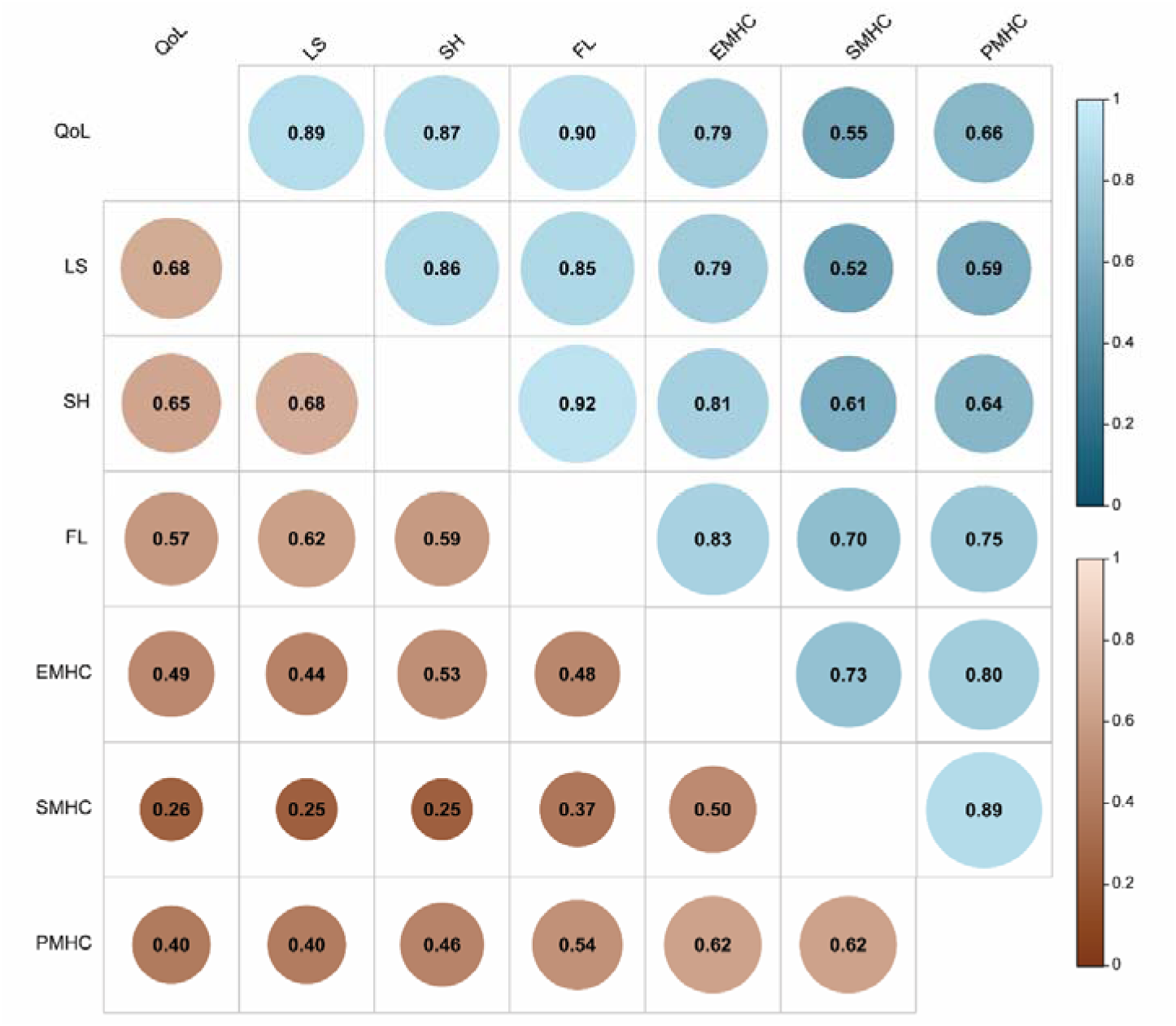
Genetic and environmental correlations Upper diagonal, in blue: Genetic correlation matrix between the wellbeing measures including the MHC as the three subscales. Bottom diagonal, in orange: Environmental correlation matrix between the wellbeing measures. QoL: Quality of Life; LS: Life Satisfaction; SH: Subjective Happiness; FL: Flourishing; EMHC: Emotional Mental Health Continuum, SMHC: Social Mental Health Continuum, PMHC: Psychological Mental Health Continuum.

The environmental correlations between the Mental Health Continuum subscales amongst each other ranged from 0.50 [95% CI = 0.44-0.55] to 0.62 [95% CI = 0.58-0.66]. The environmental correlations between the Mental Health Continuum subscales and the other wellbeing measures ranged between 0.25 [95% CI = 0.18-0.31] to 0.54 [95% CI = 0.49-0.58]; while among the other wellbeing measures, it ranged from 0.57 [95% CI = 0.52-0.61] to 0.68 [95% CI = 0.65-0.72] (Figure 4).

## Discussion

The objective of this research was to shed light on the genetic architecture of the Mental Health Continuum and its associations with other wellbeing measures. For this aim, we fitted different multivariate twin models. We showed that, genetically, the Mental Health Continuum is best explained by the three distinct subscales instead of a latent factor, hereby rendering the creation of an overall, averaged score suboptimal. When considering the Mental Health Continuum together with the other wellbeing measures, i.e., quality of life, life satisfaction, subjective happiness and flourishing, we found moderate to high genetic correlations (r = 0.52 – 0.83) between these MHC subscales and the other wellbeing measures. However, a common genetic factor underlying all the measures as previously found for a subset of these measures (Bartels & Boomsma, 2009) was not applicable.

### Genetic model of the Mental Health Continuum

We aimed to test whether, at the genetic level, the Mental Health Continuum was better explained by three subscales, i.e., emotional, social, and psychological, or by a common underlying factor estimated by a latent common factor. This last model would support the scale manual guidelines recommending averaging the items into a total score. Nevertheless, the best fitting model includes the three subscales separately (7-variable AE model), and therefore, neither the latent factor (common pathway AE model) nor the Mental Health Continuum total score (5-variable AE model) were supported.

At the same time, from the saturated model it was clear that phenotypically, the correlation between the Mental Health Continuum total score and quality of life, life satisfaction, and subjective happiness did not differ significantly from the emotional and psychological subscales, with an exception for the social subscale. This implies that when considering the Mental Health Continuum at the phenotypic level as one overall scale, we may lose trait/scale specific information about emotional, social, and psychological wellbeing on other biological layers, like genetics. Regarding the phenotypic correlations with flourishing, there were no significant differences between the Mental Health Continuum total score and the subscales. Thus, when the Mental Health Continuum is considered phenotypically, the total score or the three subscales do not show large difference in their associations with other wellbeing measures, but when considered genetically, the three subscales are recommended over the total score or a latent common factor. This implies that the MHC captures distinct underlying genetic influences that are masked when the total score is used. Consequently, studies interested in the etiology or heritability of wellbeing may lose valuable information when analyzing MHC as one construct. The results therefore highlight the importance of aligning the level of measurement (phenotypic vs. genetic) with the research question.

### Shared genetic architecture of quality of life, life satisfaction, subjective happiness, flourishing and the Mental Health Continuum

After establishing the genetic architecture of the Mental Health Continuum, we investigated the underlying sources of the relations between the wellbeing measures. The phenotypic correlations between the Mental Health Continuum and quality of life and subjective happiness are 0.52 for quality of life and 0.56 for subjective happiness, which we cannot compare with previous studies given the lack of published studies. Our correlation (0.64) between the Mental Health Continuum and flourishing is in line with the previously reported ∼0.65 for flourishing (Ploke et al., 2024). The correlation between the Mental Health Continuum and life satisfaction, however, is higher than previously reported: 0.50 in our study vs the previously reported ∼0.37 for life satisfaction (Keyes et al., 2008). This could be because of our larger sample size (*N =* 5,212) compared to theirs (*N =* 1,050). The fact that the correlation of the Mental Health Continuum total score with flourishing (0.64) are larger than with the other three measures (0.52 for quality of life, 0.50 for life satisfaction and 0.56 for subjective happiness) is also in line with previous work (Keyes, 2002; Westerhof & Keyes, 2010). This indicates that the Mental Health Continuum captures more eudaimonic aspects of wellbeing even though it was designed to capture overall wellbeing (Keyes, 2002; Westerhof & Keyes, 2010).

With respect to genetic relations between the Mental Health Continuum subscales and the other wellbeing measures, the social and psychological scales seem to be more correlated with flourishing (eudaimonic wellbeing) compared to quality of life, life satisfaction and subjective happiness. The emotional subscale seems to capture mostly both eudaimonic (flourishing) and hedonic (quality of life, life satisfaction and happiness) wellbeing since it correlates equally strongly (and the strongest) with all other measures, phenotypically (*r* = 0.56-0.62), environmentally (*rE* = 0.44-0.53), and genetically (*rG* = 0.79-0.83). For the social subscale, the gap between the phenotypic correlations with flourishing (*r* = 0.50) and the other three more hedonic measures (*r* = 0.36-0.39) is largest, driven by higher genetic and environmental correlations. The social subscale thus seems to be mostly tapping into eudaimonic wellbeing. The psychological subscale is somewhat in between the other two subscales; the gap in phenotypic correlations with flourishing and the others is present, but smaller. Once again, this is also seen in the genetic and environmental correlations between the psychological subscale and quality of life (*rG* = 0.66; *rE* = 0.40), life satisfaction (*rG* = 0.59; *rE* = 0.40), subjective happiness (*rG* = 0.64; *rE* = 0.46), and flourishing (*rG* = 0.75; *rE* = 0.54).

Taken together, these results answer an important question: are the different measures of wellbeing interchangeable, or do they cover different nuances of wellbeing? The genetic correlations (Figure 4) among the wellbeing measures were moderate to high, suggesting substantial overlap in the underlying liabilities. In previous studies, it was shown that the wellbeing measures quality of life, life satisfaction, and subjective happiness were influenced by a common genetic factor (Bartels & Boomsma, 2009). However, when an independent pathway model was tested in our data, the genetic and environment components could not be constrained to one underlying factor. We thus did not observe a clear common genetic and/or environmental factor underlying all the measures.

In the case of a GWAS, where a large sample size is required, the genetic correlations found here are high enough to consider the distinct wellbeing measures interchangeably to increase the sample size using multivariate GWAS techniques (Ruotsalainen et al., 2021) - albeit at the cost of increasing heterogeneity in the phenotype, potentially deflating SNP-based heritability estimates (Manchia et al., 2013). If however, one is more interested in the distinct determinants of the specific dimensions of wellbeing, such as emotional, social and psychological, or even hedonic and eudaimonic, it may not be ideal to treat the different measures as one, since there are observed differences in the genetics of the subconstructs.

In an era in which big data is essential for research, the findings of this study can facilitate future collaborations across cohorts, both at the phenotypic and genetic level. Because cohorts use a wide range of wellbeing measures, understanding how these measures relate to one another is critical. The better our understanding of the different wellbeing measures, the easier it will be to facilitate variable harmonization and collaborations, for example for meta-analyses, with other cohorts having different wellbeing measurements available.

### Strengths and limitations

One of the strengths of this study is the inclusion of both hedonic and eudaimonic measures, allowing us to capture a broad range of individual differences in wellbeing. Another important strength is the large overall sample size (*N* = 5,212). Despite this advantage, the number of complete twin pairs available for analysis was relatively limited (*N* = 1,024). In addition, the Netherlands Twin Register (NTR) sample is overrepresented by women and participants with a relatively high level of educational attainment, resulting in a sex distribution that is not balanced (3750 females/1462 males) and a cohort that is not fully representative of the general population since it is of a higher social economic status. These characteristics may limit the generalizability of the findings. Finally, we note the inherent limitations associated with twin modelling approaches, including assumptions such as the absence of assortative mating, the equal environments assumption, and the fact that standard twin models assume no gene-environment correlation and interaction (Polderman et al., 2015). Results should be interpreted in light of these limitations.

### Conclusion

Overall, we showed that the Mental Health Continuum is best described, genetically, as three separate but correlated subscales, capturing emotional, social and psychological wellbeing. Among these, the emotional scale shows the strongest genetic correlation with the other wellbeing measures, both hedonic (quality of life, life satisfaction and subjective happiness) and eudaimonic (flourishing).The results show that in wellbeing research for large-scale phenotypic research, relatively broad measures, such as quality of life, may be sufficient from a pragmatic perspective. However, for research that is more genetically or mechanistically oriented, more targeted and conceptually specific measures of wellbeing may be required.

## Statements and Declarations

The authors have no competing interests to declare that are relevant to the content of this article. This study is funded by an NWO-VICI grant (VI.C.211.054, PI Bartels). Data collection for this study has been funded by ERC Consolidator Grant (WELL-BEING; grant 771057).

## Acknowledgements

We warmly thank all participating twins in the Netherlands Twin Register who dedicated part of their time to make research possible as well as everyone involved in the collection of the data and data management.

## Notes

### Competing Interest Statement

The authors have declared no competing interest.

