## Supplementary Material for "The multidimensional structure of wellbeing: genetic evidence from a multivariate twin study including the Mental Health Continuum"

Behavior Genetics

Supplementary Table 1. Age and Sex effects on the saturated model

| base | comparison | Estimated parameters | -2LL(1) | df(2) | AIC(3) | ΔLL(4) | Δdf(5) | P(6) |
|---|---|---|---|---|---|---|---|---|
| *Saturated* | - | 231 | 77329.40 | 36253 | 77791.40 | NA | NA | NA |
| *No age effects* | Saturated | 224 | 77380.94 | 36260 | 77828.94 | 51.54 | 7 | 0 |
| *No sex effects* | Saturated | 224 | 77370.48 | 36260 | 77818.48 | 41.07 | 7 | 0 |

Table 1. Note: 1. The negative log-likelihood subtracted for the more general model from the -2LL of the more restricted model. 2. Degrees of freedom (df) of the model. 3. The Akaike information criterion (AIC) is an estimator of prediction error and thereby relative quality of statistical models for a given set of data. 4. $X^2$ test given in the −2LL test. 5. Difference of degrees of freedom equal to the difference in the number of estimated parameters in the two models. 6. *P* represents the *p*-value of the test performed. In bold are the models with no difference from the original model; therefore, the best fitting models.

Supplementary Table 2. Constrains saturated model

Means across twin order and zygosity were first constrained in the saturated model (Supplementary Table 1). No twin order effect on variances was observed ($P_{FDR}$ = .33). Within-person cross-trait covariances could be constrained across twin order ($P_{FDR}$ = .15), but not the cross-twin cross-trait covariances ($P_{FDR}$ = .00). However, the differences in correlations were small (for MZ twins, the median of the absolute differences was 0.03 (SD = 0.03), and for DZ twins, it was 0.03 (SD = 0.03)). Therefore, even though the Chi-square difference test was significant, the differences appeared to be negligible in an absolute matter. We, therefore, chose to continue with the more parsimonious, constrained model. Variances could not be constrained for MZ twins and DZ twins ($P_{FDR}$ = .00). However, again, the median of the absolute differences between the MZ and DZ variances was 0.04 (SD = 0.03), and we proceeded as such with the constrained model. For all 5 measures, within-person cross-trait covariances were constrained for MZ twins and DZ twins ($P_{FDR}$ = .33).

| base | comparison | Estimated parameters | -2LL(1) | df(2) | AIC(3) | ΔLL(4) | Δdf(5) | P(6) |
|---|---|---|---|---|---|---|---|---|

| | | | | | | | | |
|---|---|---|---|---|---|---|---|---|
| *Saturated* | - | 231 | 77329.40 | 36253 | 77791.40 | NA | NA | NA |
| *Equal means and variances across twin order* | Saturated | 217 | 77345.65 | 36267 | 77779.65 | 16.24 | 14 | 0.33 |
| *Equal covariances across twin order* | *Equal means and variances across twin order* | 175 | 77398.71 | 36309 | 77748.71 | 53.07 | 42 | 0.15 |
| *Equal covariances Twin1-Twin2 vs Twin2-Twin1* | *Equal covariances across twin order* | 133 | 77472.62 | 36351 | 77738.62 | 73.90 | 42 | 0.00 |
| *Equal variances across zygosity* | *Equal covariances Twin1-Twin2 vs Twin2-Twin1* | 126 | 77495.37 | 36358 | 77747.37 | 22.76 | 7 | 0.00 |
| *Equal covariances across zygosity* | *Equal variances across zygosity* | 105 | 77519.56 | 36379 | 77729.56 | 24.19 | 21 | 0.33 |

Supplementary Table 2. Note: 1. The negative log-likelihood subtracted for the more general model from the -2LL of the more restricted model. 2. Degrees of freedom (df) of the model. 3. The Akaike information criterion (AIC) is an estimator of prediction error and thereby relative quality of statistical models for a given set of data. 4. $X^2$ test given in the −2LL test. 5. Difference of degrees of freedom equal to the difference in the number of estimated parameters in the two models. 6. *P* represents the *p*-value of the test performed.

Supplementary Table 3. Comparison within-model

| base | comparison | Estimated parameters | -2LL(1) | df(2) | AIC(3) | ΔLL(4) | Δdf(5) | P(6) |
|---|---|---|---|---|---|---|---|---|
| *ACE_5v* | - | 58 | 56063.33 | 26022 | 56179.33 | NA | NA | NA |
| ***AE_5v*** | **ACE_5v** | **43** | **56082.85** | **26017** | **56167.85** | **19.53** | **15** | **0.19** |
| *CE_5v* | ACE_5v | 43 | 56117.44 | 26017 | 56203.44 | 54.11 | 15 | 2.51e-06 |
| *E_5v* | ACE_5v | 28 | 56415.32 | 26032 | 56471.32 | 351.99 | 30 | 2.26e-56 |
| *ACE_7v* | - | 102 | 77214.69 | 36382 | 77418.69 | NA | NA | NA |
| ***AE_7v*** | **ACE_7v** | **74** | **77253.62** | **36410** | **77401.62** | **38.93** | **28** | **0.08** |
| *CE_7v* | ACE_7v | 74 | 77293.79 | 36410 | 77441.79 | 79.10 | 28 | 9.07e-07 |
| *E_7v* | ACE_7v | 46 | 77658.64 | 36438 | 77750.64 | 443.95 | 56 | 9.25e-62 |
| *ACE_cP* | - | 74 | 78092.53 | 36410 | 78240.53 | NA | NA | NA |
| ***AE_cP*** | **ACE_cP** | **56** | **78116.94** | **36428** | **78228.94** | **24.41** | **18** | **0.14** |
| *CE_cP* | ACE_cP | 59 | 78142.27 | 36425 | 78260.27 | 49.74 | 15 | 1.33e-05 |
| *E_cP* | ACE_cP | 41 | 78417.54 | 36443 | 78499.54 | 325.01 | 33 | 1.05e-49 |
| *ACE_iP* | - | 60 | 77936.79 | 36424 | 78056.79 | NA | NA | NA |
| *AE_iP* | ACE_iP | 46 | 78125.55 | 36438 | 78217.55 | 188.76 | 14 | 1.08e-32 |

Supplementary Table 3. Note: 1. The negative log-likelihood subtracted for the more general model from the -2LL of the more restricted model. 2. Degrees of freedom (df) of the model. 3. The Akaike information criterion (AIC) is an estimator of prediction error and thereby relative quality of statistical models for a given set of data. 4. $X^2$ test given in the −2LL test. 5. Difference of degrees of freedom equal to the difference in the number of estimated parameters in the two models. 6. *P* represents the *p*-value of the test performed.

Supplementary Table 4. Unstandardized variances AE_7v model

| | QoL | LS | SH | FL | EMHC | SMHC | PMHC |
|---|---|---|---|---|---|---|---|
| QoL | 0.38 / 0.61 | 0.35 | 0.35 | 0.35 | 0.27 | 0.22 | 0.25 |
| LS | 0.41 | 0.42 / 0.59 | 0.36 | 0.34 | 0.28 | 0.21 | 0.23 |
| SH | 0.39 | 0.40 | 0.41 / 0.59 | 0.37 | 0.29 | 0.24 | 0.25 |
| FL | 0.35 | 0.37 | 0.35 | 0.39 / 0.62 | 0.28 | 0.28 | 0.28 |
| EMHC | 0.31 | 0.28 | 0.33 | 0.31 | 0.30 / 0.69 | 0.25 | 0.27 |
| SMHC | 0.16 | 0.15 | 0.15 | 0.23 | 0.32 | 0.40 / 0.60 | 0.34 |
| PMHC | 0.25 | 0.28 | 0.28 | 0.33 | 0.41 | 0.38 | 0.37 / 0.63 |

Supplementary Table 4. Upper diagonal A unstandardized variances. Bottom diagonal E unstandardized variances.

Supplementary Table 5. Summary estimates AE_5v model

| name | matrix | row | col | Estimate | Std.Error |
|---|---|---|---|---|---|
| VA11 | VA | 1 | 1 | 0.38208144 | 0.03203726 |
| VA21 | VA | 1 | 2 | 0.3548671 | 0.0277734 |
| VA22 | VA | 2 | 2 | 0.4146394 | 0.03162364 |
| VA31 | VA | 1 | 3 | 0.34595669 | 0.02783424 |
| VA32 | VA | 2 | 3 | 0.35514494 | 0.02783793 |
| VA33 | VA | 3 | 3 | 0.41069804 | 0.03220152 |
| VA41 | VA | 1 | 4 | 0.34799025 | 0.02713612 |
| VA42 | VA | 2 | 4 | 0.34240333 | 0.02727934 |
| VA43 | VA | 3 | 4 | 0.3679624 | 0.02756437 |
| VA44 | VA | 4 | 4 | 0.39066849 | 0.03240561 |
| VA51 | VA | 1 | 5 | 0.27067631 | 0.02544585 |
| VA52 | VA | 2 | 5 | 0.26320486 | 0.02492853 |
| VA53 | VA | 3 | 5 | 0.28832989 | 0.0258739 |
| VA54 | VA | 4 | 5 | 0.31944404 | 0.02670579 |
| VA55 | VA | 5 | 5 | 0.41997334 | 0.03226426 |
| VE11 | VE | 1 | 1 | 0.60988052 | 0.02964014 |
| VE21 | VE | 1 | 2 | 0.40491122 | 0.02496893 |
| VE22 | VE | 2 | 2 | 0.59145212 | 0.02870332 |
| VE31 | VE | 1 | 3 | 0.38711109 | 0.02499447 |
| VE32 | VE | 2 | 3 | 0.40226834 | 0.02482927 |

| | | | | | |
|---|---|---|---|---|---|
| VE33 | VE | 3 | 3 | 0.585529 | 0.02916346 |
| VE41 | VE | 1 | 4 | 0.34954282 | 0.02439764 |
| VE42 | VE | 2 | 4 | 0.3743381 | 0.02448656 |
| VE43 | VE | 3 | 4 | 0.35264518 | 0.02448808 |
| VE44 | VE | 4 | 4 | 0.6159203 | 0.02993354 |
| VE51 | VE | 1 | 5 | 0.25357322 | 0.02280898 |
| VE52 | VE | 2 | 5 | 0.24206915 | 0.02216558 |
| VE53 | VE | 3 | 5 | 0.26971305 | 0.022943 |
| VE54 | VE | 4 | 5 | 0.32177897 | 0.02384695 |
| VE55 | VE | 5 | 5 | 0.58254219 | 0.02904795 |
| meanQoL | MZ.meanG | 1 | 1 | -0.3234662 | 0.04064121 |
| meanSat | MZ.meanG | 1 | 2 | -0.0506802 | 0.04089734 |
| meanHap | MZ.meanG | 1 | 3 | -0.2875493 | 0.03806779 |
| meanFlo | MZ.meanG | 1 | 4 | -0.0173539 | 0.03814916 |
| meanMHC | MZ.meanG | 1 | 5 | -0.0743556 | 0.04236291 |
| bAge_QoL | MZ.b1 | 1 | 1 | 0.98055107 | 0.09700858 |
| bAge_Sat | MZ.b1 | 1 | 2 | 0.23317288 | 0.09811669 |
| bAge_Hap | MZ.b1 | 1 | 3 | 0.79187059 | 0.09756232 |
| bAge_Flo | MZ.b1 | 1 | 4 | 0.06918263 | 0.09772559 |
| bAge_MHC | MZ.b1 | 1 | 5 | 0.18226648 | 0.09803906 |
| bSex_QoL | MZ.b2 | 1 | 1 | -0.0437037 | 0.01942518 |
| bSex_Sat | MZ.b2 | 1 | 2 | -0.0470073 | 0.01896411 |
| bSex_MHC | MZ.b2 | 1 | 5 | 0.00580757 | 0.02405464 |

Supplementary Table 6. Summary estimates AE_7v model

| name | matrix | row | col | Estimate | Std.Error |
|---|---|---|---|---|---|
| VA11 | VA | 1 | 1 | 0.38193828 | 0.03201631 |
| VA21 | VA | 1 | 2 | 0.35449465 | 0.02773094 |
| VA22 | VA | 2 | 2 | 0.41593508 | 0.03154048 |
| VA31 | VA | 1 | 3 | 0.34581768 | 0.02777544 |
| VA32 | VA | 2 | 3 | 0.35530148 | 0.02774971 |
| VA33 | VA | 3 | 3 | 0.41050664 | 0.0320936 |
| VA41 | VA | 1 | 4 | 0.34748811 | 0.02710449 |
| VA42 | VA | 2 | 4 | 0.34246368 | 0.02722874 |
| VA43 | VA | 3 | 4 | 0.36756342 | 0.02748802 |
| VA44 | VA | 4 | 4 | 0.38883021 | 0.03236998 |
| VA51 | VA | 1 | 5 | 0.26800859 | 0.0270292 |
| VA52 | VA | 2 | 5 | 0.28009576 | 0.02649405 |
| VA53 | VA | 3 | 5 | 0.2865788 | 0.02747392 |
| VA54 | VA | 4 | 5 | 0.28487081 | 0.02729837 |
| VA55 | VA | 5 | 5 | 0.30379096 | 0.035105 |
| VA61 | VA | 1 | 6 | 0.21543345 | 0.02395853 |
| VA62 | VA | 2 | 6 | 0.21199772 | 0.02360268 |

| | | | | | |
|---|---|---|---|---|---|
| VA63 | VA | 3 | 6 | 0.24464801 | 0.02405576 |
| VA64 | VA | 4 | 6 | 0.2751674 | 0.02501989 |
| VA65 | VA | 5 | 6 | 0.25279689 | 0.02721167 |
| VA66 | VA | 6 | 6 | 0.39747339 | 0.0320063 |
| VA71 | VA | 1 | 7 | 0.24733164 | 0.02532993 |
| VA72 | VA | 2 | 7 | 0.2295759 | 0.02479783 |
| VA73 | VA | 3 | 7 | 0.24991681 | 0.02576893 |
| VA74 | VA | 4 | 7 | 0.28476648 | 0.02668326 |
| VA75 | VA | 5 | 7 | 0.26778981 | 0.02891577 |
| VA76 | VA | 6 | 7 | 0.34016225 | 0.02775792 |
| VA77 | VA | 7 | 7 | 0.37015422 | 0.03260327 |
| VE11 | VE | 1 | 1 | 0.60986334 | 0.02963356 |
| VE21 | VE | 1 | 2 | 0.40517147 | 0.02492849 |
| VE22 | VE | 2 | 2 | 0.59027969 | 0.02858771 |
| VE31 | VE | 1 | 3 | 0.38713334 | 0.02495613 |
| VE32 | VE | 2 | 3 | 0.40213786 | 0.02474259 |
| VE33 | VE | 3 | 3 | 0.58574986 | 0.02908573 |
| VE41 | VE | 1 | 4 | 0.3499965 | 0.02439701 |
| VE42 | VE | 2 | 4 | 0.37430759 | 0.02444441 |
| VE43 | VE | 3 | 4 | 0.3530244 | 0.02444724 |
| VE44 | VE | 4 | 4 | 0.61763953 | 0.02995955 |
| VE51 | VE | 1 | 5 | 0.31457444 | 0.02515767 |
| VE52 | VE | 2 | 5 | 0.2811152 | 0.02436168 |
| VE53 | VE | 3 | 5 | 0.33432134 | 0.02531479 |
| VE54 | VE | 4 | 5 | 0.31179773 | 0.02532025 |
| VE55 | VE | 5 | 5 | 0.68756373 | 0.03399593 |
| VE61 | VE | 1 | 6 | 0.15559993 | 0.02170144 |
| VE62 | VE | 2 | 6 | 0.14673855 | 0.02117253 |
| VE63 | VE | 3 | 6 | 0.1457738 | 0.02145378 |
| VE64 | VE | 4 | 6 | 0.22516304 | 0.02250604 |
| VE65 | VE | 5 | 6 | 0.31836054 | 0.02530333 |
| VE66 | VE | 6 | 6 | 0.5990606 | 0.02930435 |
| VE71 | VE | 1 | 7 | 0.24657306 | 0.023127 |
| VE72 | VE | 2 | 7 | 0.24557113 | 0.02251684 |
| VE73 | VE | 3 | 7 | 0.27908863 | 0.0233744 |
| VE74 | VE | 4 | 7 | 0.33484973 | 0.02437754 |
| VE75 | VE | 5 | 7 | 0.41012033 | 0.02718968 |
| VE76 | VE | 6 | 7 | 0.38122823 | 0.02509956 |
| VE77 | VE | 7 | 7 | 0.62875698 | 0.03041541 |
| meanQoL | MZ.meanG | 1 | 1 | -0.3174533 | 0.04025179 |
| meanSat | MZ.meanG | 1 | 2 | -0.0488121 | 0.04052102 |
| meanHap | MZ.meanG | 1 | 3 | -0.2886952 | 0.03772718 |
| meanFlo | MZ.meanG | 1 | 4 | -0.0179991 | 0.03768454 |
| meanEMHC | MZ.meanG | 1 | 5 | -0.2438268 | 0.03720689 |
| meanSMHC | MZ.meanG | 1 | 6 | -0.0470826 | 0.04195017 |
| meanPMHC | MZ.meanG | 1 | 7 | 0.09942082 | 0.04083227 |

| | | | | | |
|---|---|---|---|---|---|
| bAge_QoL | MZ.b1 | 1 | 1 | 0.97937532 | 0.09604466 |
| bAge_Sat | MZ.b1 | 1 | 2 | 0.23442308 | 0.0971036 |
| bAge_Hap | MZ.b1 | 1 | 3 | 0.79394662 | 0.09660763 |
| bAge_Flo | MZ.b1 | 1 | 4 | 0.06981463 | 0.09643151 |
| bAge_EMHC | MZ.b1 | 1 | 5 | 0.65807292 | 0.09507106 |
| bAge_SMHC | MZ.b1 | 1 | 6 | 0.23298444 | 0.09675547 |
| bAge_PMHC | MZ.b1 | 1 | 7 | -0.1658086 | 0.09636887 |
| bSex_QoL | MZ.b2 | 1 | 1 | -0.0517326 | 0.01919414 |
| bSex_Sat | MZ.b2 | 1 | 2 | -0.0506118 | 0.01889915 |
| bSex_SMHC | MZ.b2 | 1 | 6 | -0.053761 | 0.02407258 |
| bSexPMHC | MZ.b2 | 1 | 7 | -0.0586407 | 0.02079563 |
